# The ER cargo receptor SURF4 facilitates efficient erythropoietin secretion

**DOI:** 10.1101/866954

**Authors:** Zesen Lin, Richard King, Vi Tang, Greggory Myers, Ginette Balbin-Cuesta, Ann Friedman, Beth McGee, Karl Desch, Ayse Bilge Ozel, David Siemieniak, Pavan Reddy, Brian Emmer, Rami Khoriaty

## Abstract

Erythropoietin (EPO), a glycoprotein produced by specialized peritubular fibroblasts in the kidney, is the master regulator of erythropoiesis. EPO is secreted into the plasma in response to tissue hypoxia and stimulates erythroid differentiation and maturation. Though the transcriptional regulation of EPO has been well studied, the molecular determinants of EPO secretion remain unknown. Here, we generated a HEK293T reporter cell line that provides a quantifiable and selectable readout of intracellular EPO levels. Using this cell line, we performed a genome-scale CRISPR screen that identified SURF4 as an important mediator of EPO secretion. Targeting *SURF4* with multiple independent sgRNAs resulted in intracellular accumulation and extracellular depletion of EPO. Both of these phenotypes were rescued by expression of *SURF4* cDNA. Additionally, consistent with a role for SURF4 as an ER cargo receptor of EPO, we found that disruption of SURF4 resulted in accumulation of EPO in the ER compartment, and that SURF4 and EPO physically interact. Furthermore, SURF4 disruption in Hep3B cells also caused a defect in the secretion of endogenous EPO, ruling out an artifact of heterologous overexpression. This work suggests that SURF4 functions as an ER cargo receptor that mediates the efficient secretion of EPO. Our findings also suggest that modulating SURF4 may be an effective treatment for disorders of erythropoeisis that are driven by aberrant EPO levels. Finally, we show that SURF4 overexpression results in increased secretion of EPO, suggesting a new strategy for more efficient production of recombinant EPO.

## Introduction

Approximately one third of the proteins encoded by the mammalian genome are secretory proteins (1, 2). These proteins traffic from the endoplasmic reticulum (ER) to the Golgi apparatus via coat protein complex II (COPII) vesicles before reaching their final destinations: endosomes, lysosomes, plasma membrane, or extracellular space. COPII vesicles have an inner coat composed of SAR1 and SEC23-SEC24 heterodimers and an outer coat composed of SEC13-SEC31 heterotetramers (3). Though transmembrane cargo proteins may directly interact with COPII components, the physical barrier created by the ER membrane prevents direct interaction between soluble cargos and the COPII coat. Therefore, soluble cargos either passively flow into COPII vesicles (bulk flow) or are captured in COPII vesicles through recognition by intermediary receptors or adaptors (cargo capture) (4).

Support for receptor-mediated cargo capture arose from early electron microscopy studies and *in vitro* assays of cargo packaging in COPII vesicles, which demonstrated efficient selection and concentration of cargos into COPII vesicles, as well as physical interactions between soluble cargos and COPII components (4–9). Subsequent studies uncovered LMAN1 as the first ER cargo receptor that mediates ER export of soluble cargos in mammals (10–12). LMAN1, together with its adapter MCFD2, form a complex that is required for the efficient secretion of coagulation factors V and VIII, cathepsins, and alpha1-antitrypsin (12–16). While a handful of additional interactions between soluble cargos and ER receptors have been described in mammals (4, 9, 17), the extent to which bulk flow and cargo capture contribute to recruitment of proteins in COPII vesicles is unclear. This is primarily due to the fact that ER cargo receptors that are necessary for the efficient secretion of the majority of soluble cargos remain unidentified.

Erythropoietin (EPO) is a glycoprotein that is produced predominantly by specialized kidney peritubular fibroblasts and secreted into the plasma (18–21). EPO binds to its receptor expressed on erythroid precursors and promotes cell survival, proliferation, and differentiation (22–24). EPO plays a crucial role in the regulation of red blood cell production (erythropoiesis). Failure to make sufficient amounts of EPO, as seen in the setting of chronic kidney disease, results in anemia. In contrast, supra-physiological EPO levels, as seen in the context of EPO-secreting tumors, result in polycythemia. Though the transcriptional regulation of EPO production has been well-studied (25–30), the molecular basis of EPO trafficking remains poorly understood.

In this study, in an effort to identify proteins involved in EPO secretion, we developed a genome-scale CRISPR/Cas9 knock-out screen that provides a quantifiable and selectable readout of intracellular EPO levels. This screen, followed by several validation experiments, identified the ER cargo receptor SURF4 as a key mediator of efficient EPO secretion. These findings suggest that modulation of SURF4 in the EPO producing cells could provide a novel strategy for altering plasma EPO levels and therefore regulating erythropoiesis. Additionally, this report suggests a novel strategy for improving the efficiency of recombinant EPO production.

## Results

### Generation of a reporter cell line that allows for a quantifiable and selectable readout of intracellular EPO levels

To identify proteins that regulate the intracellular trafficking of EPO, we developed a genome-scale functional screen that provides a quantifiable and selectable readout of intracellular EPO accumulation. Therefore, we generated a reporter HEK293T cell line stably expressing eGFP-tagged EPO (EPO-eGFP) and, as an internal control, mCherry-tagged alpha1-antitrypsin (A1AT-mCherry) (Fig. 1*A*). This cell line is herein referred to as the EPO-eGFP/A1AT-mCherry reporter cell line or just as the reporter cell line. Importantly, EPO-eGFP and A1AT-mCherry are equally expressed from the same CMV promoter, with their respective coding sequences separated by a sequence encoding a P2A peptide (Fig. 1*A*).

We found that both EPO and A1AT are efficiently secreted from the reporter cell line (Fig. 1 *B* and *C*) and that disruption of ER-to-Golgi transport with Brefeldin A results in intracellular accumulation of EPO and A1AT (Fig. 1*D*). Deletion of the ER cargo receptor for A1AT, *LMAN1*, resulted in intracellular accumulation of A1AT, as expected, with no effect on intracellular EPO (Fig. 1*E*), ruling out a role for LMAN1 in EPO secretion. These studies demonstrate that the machinery required for the efficient secretion of EPO via the classical secretory pathway is intact in this cell line and establish the utility of this cell line to identify modifiers of intracellular EPO levels.

**Fig. 1.**
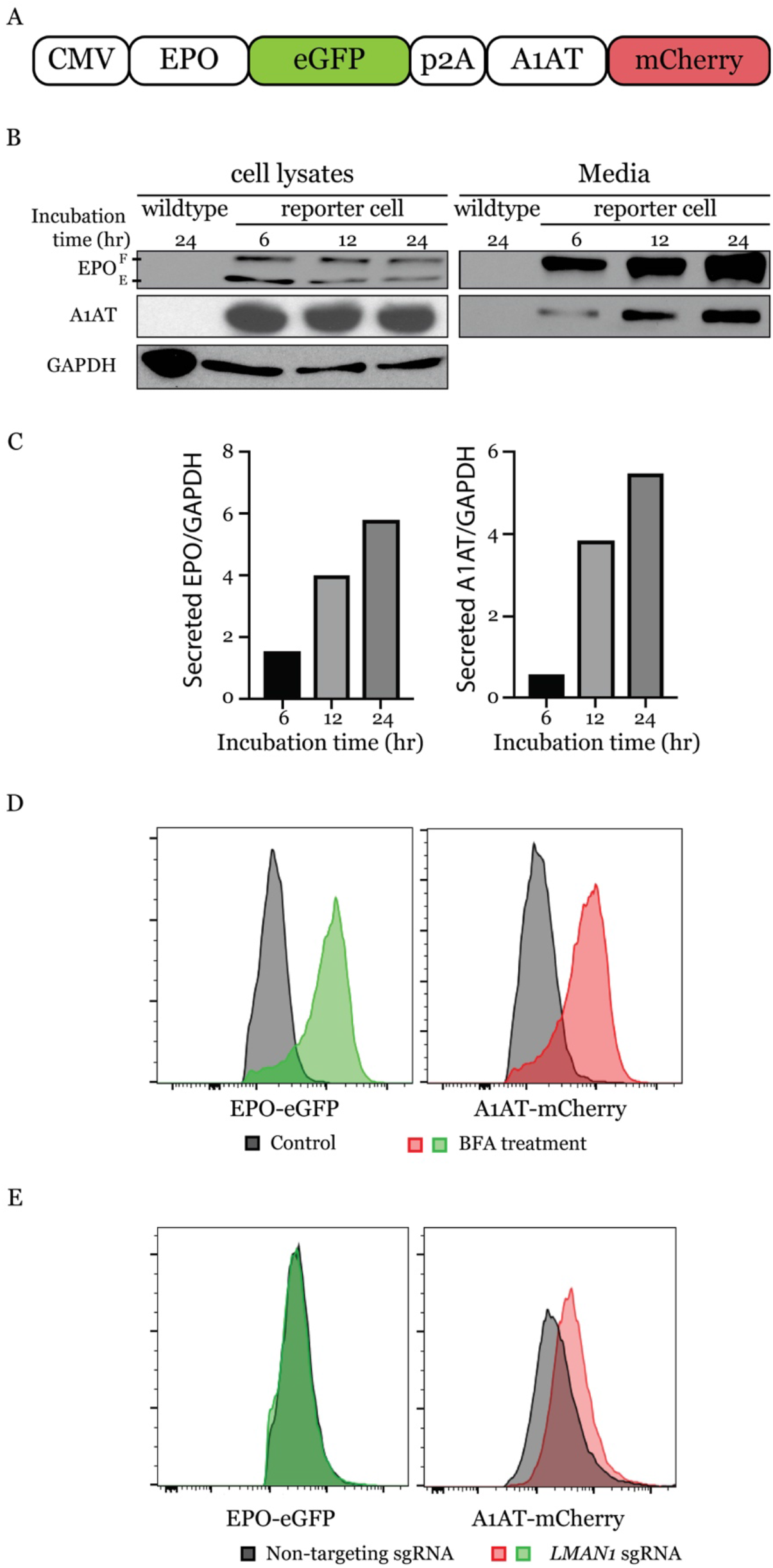
A reporter HEK293T cell line stably expressing EPO-eGFP and A1AT-mCherry. (*A*) A construct that expresses EPO-eGFP and A1AT-mCherry from the same CMV promoter was assembled and used to generate the reporter cell line. A P2A sequence separates EPO-eGFP from A1AT-mCherry. (*B*) Intracellular and extracellular EPO-eGFP and A1AT-mCherry protein abundance was determined by Western blot using anti-eGFP and anti-mCherry antibodies, respectively. E: ER form of EPO; F: fully-glycosylated EPO. (*C*) Protein abundance was quantified using Image J with GAPDH as control. (*D*) Inhibiting ER to Golgi transport with Brefeldin A (BFA) leads to intracellular accumulation of EPO-eGFP and A1AT-mCherry, as measured by fluorescence intensity (*E*) *LMAN1* deletion results in intracellular accumulation of A1AT with no effect on EPO.

### A CRISPR/Cas9 loss-of-function screen identified SURF4 as a putative regulator of intracellular EPO level

To identify proteins that affect EPO secretion, we mutagenized the EPO-eGFP/A1AT-mCherry reporter cell line with a CRISPR/Cas9 knockout library (hGeCKO-v2), which delivers SpCas9, a puromycin resistance cassette, and a pooled collection of 123,411 single guide RNAs (sgRNAs) that include 6 sgRNAs targeting nearly every gene in the human genome. Transduction of the library was performed at a low multiplicity of infection (MOI ~0.3), such that most infected cells receive 1 sgRNA to mutate 1 gene in the genome. Puromycin selection was applied from days 1-4 post-transduction. After an additional 9 days, cells with normal mCherry but increased (top ~7%) or decreased (bottom ~7%) eGFP fluorescence were isolated (Fig. 2*A*). This cell sorting strategy allows the identification of genes that affect EPO but not A1AT levels, therefore reducing the likelihood of identifying genes that affect global protein secretion. Integrated sgRNA sequences were quantified by deep sequencing and analyzed for their enrichment in the eGFP high compared to the eGFP low population.

**Fig. 2.**
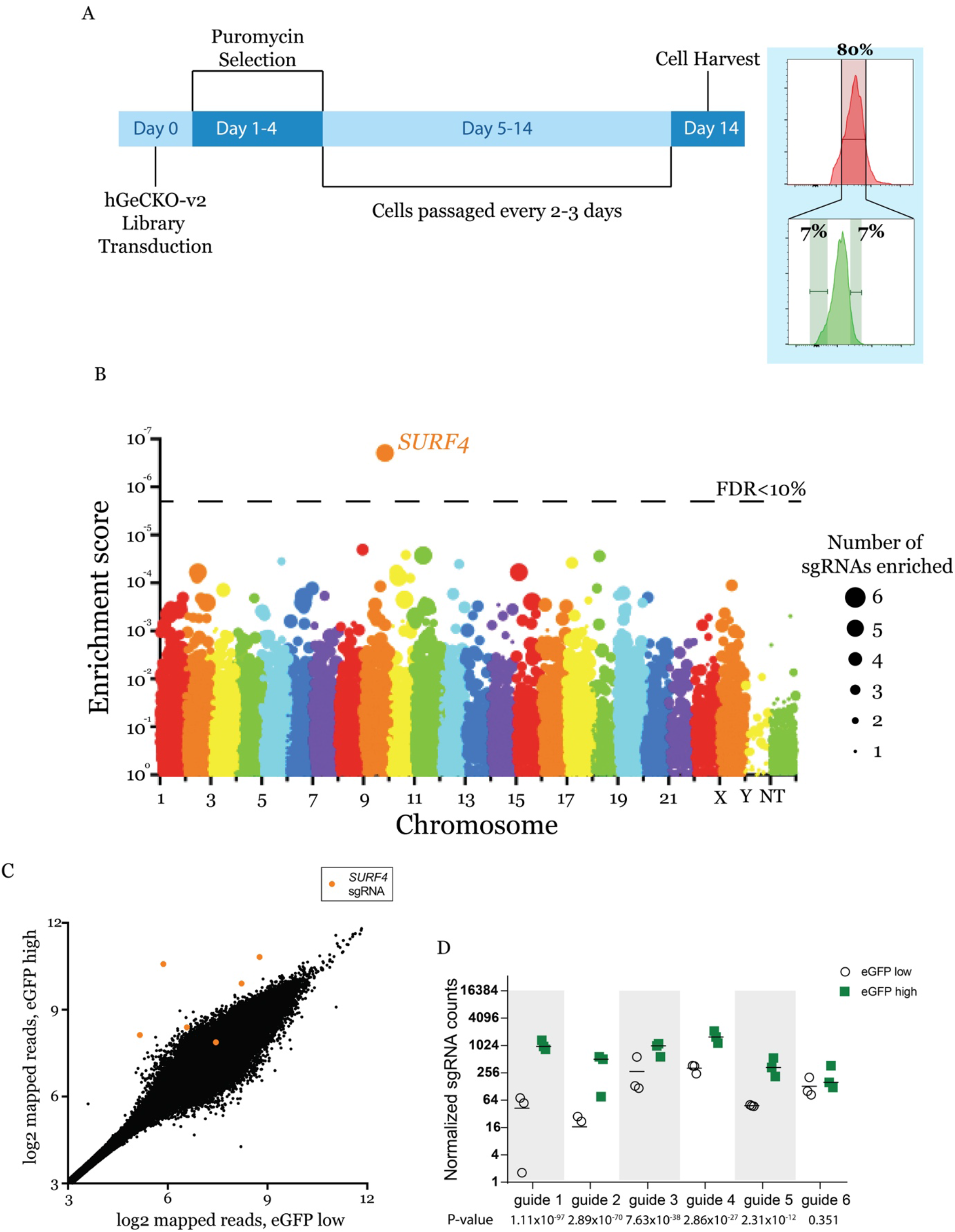
CRISPR/Cas9 loss-of-function screen to identify genes that affect intracellular EPO levels. (*A*) Screen strategy: 24 hours following transduction of the CRISPR library, puromycin selection was applied for 3 days. At day 14, cells with unchanged mCherry but with top or bottom 7% eGFP fluorescence were isolated. sgRNAs abundance was then determined in each cell population. (*B*) Gene level enrichment score was calculated for every gene using MAGeCK (see methods). Each gene is represented by a bubble, the size of which is proportional to number of sgRNAs with significant enrichment in the eGFP high population. *SURF4* has the highest MAGeCK enrichment score and is the only gene for which the false discovery rate (FDR) is <10%. NT: non-targeting. (*C*) Normalized abundance of *SURF4*-targeting sgRNAs in the eGFP high and eGFP low populations. Abundance score calculated from 3 biological replicates, using DEseq (see methods). *SURF4* sgRNAs are highlighted in orange. (*D*) Normalized counts for the 6 *SURF4* targeting sgRNAs included in the library, for all 3 biological replicates. p-values were calculated using MAGeCK.

This screen, performed in biological triplicates, identified that the sgRNA sequences targeting only one gene, surfeit locus protein 4 (*SURF4*), are enriched in the eGFP high population compared to the eGFP low population at an FDR <10% (Fig. 2*B*). Five out of six sgRNAs targeting *SURF4* were significantly enriched in the eGFP high population (Fig. 2 *C* and *D*).

### *SURF4* deletion results in intracellular accumulation and reduced secretion of EPO

To validate the results of the screen, we targeted *SURF4* with one sgRNA (sgRNA1) that showed significant enrichment in the whole genome screen (Fig. 2*D*) and a second sgRNA (sgRNA2) not included in the hGeCKO-v2 library. *SURF4* mutagenesis with sgRNA1 or sgRNA2 was highly efficient, resulting in insertions or deletions (indels) in ~97% and 77% of alleles, respectively (Fig. 3*A*). Cells transduced with *SURF4* sgRNA1 or sgRNA2 exhibited increased intracellular accumulation of EPO-eGFP, with no effect on A1AT-mCherry (Fig. 3 *B* and *C*). This finding was confirmed in 3 independent EPO-eGFP/A1AT-mCherry reporter cell clones (Fig. 3*D*), ruling out an artifact unique to the clone used in the original screen.

**Fig. 3.**
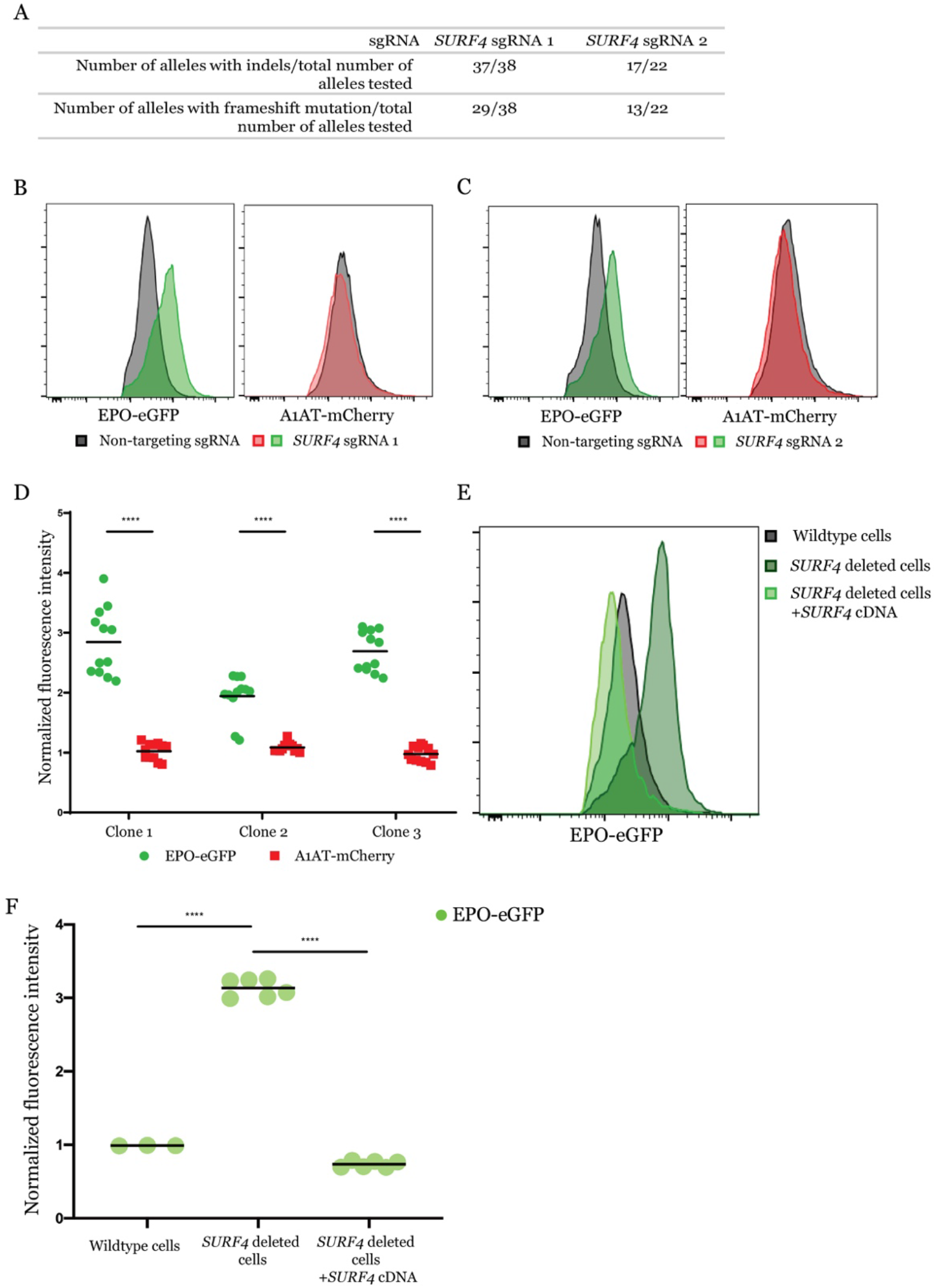
*SURF4* deletion results in intracellular accumulation of EPO-eGFP. (*A*) *SURF4*-targeting sgRNA1 and sgRNA2 are highly efficient, causing indels in ~97% and 77% of alleles, respectively. (*B, C*) Flow cytometry histograms showing intracellular accumulation of EPO, but not A1AT, following *SURF4* deletion in HEK293T cells, using 2 independent sgRNAs, (*B*) sgRNA1 or (*C*) sgRNA2. (*D*) Quantification of intracellular mean fluorescence intensity in 3 independent clonal reporter cell lines transduced with *SURF4*-sgRNA1 (n=12). Results were normalized to mean fluorescence intensity of cells transduced with non-targeting sgRNAs. (*E, F*) Flow cytometry histograms and normalized mean fluorescence intensity of EPO-eGFP in several clonal cell lines with sequence-confirmed *SURF4* frameshift mutations (*SURF4* deleted) with or without stable expression of wildtype *SURF4* cDNA. Mean fluorescence intensity in panel *F* was normalized to that of wildtype cells. **** p<0.0001.

To further confirm a direct effect of SURF4 deficiency on intracellular EPO accumulation, we next generated 3 clonal reporter cell lines with confirmed frameshift mutations of both *SURF4* alleles, by transient expression of *SURF4* sgRNA1 (*SI Appendix*, Fig. S1). The increased intracellular EPO protein levels observed in *SURF4* deleted cells was completely rescued by a lentivirus expressing wildtype *SURF4* cDNA (Fig. 3 *E* and *F*), ruling out an off-target effect shared by sgRNA1 and sgRNA2. Taken together, these findings demonstrate that *SURF4* disruption results in intracellular accumulation of EPO.

To rule out an indirect effect on EPO-eGFP secretion resulting from an interaction between eGFP and SURF4, we analyzed the dependence of FLAG-tagged EPO on SURF4 for secretion. We generated a wildtype and a SURF4 deficient HEK293 cell line expressing FLAG-tagged EPO (EPO-FLAG) from a tetracycline-inducible promoter (Fig. 4*A*). Following tetracycline administration, the intracellular EPO accumulation was significantly more pronounced in SURF4-deficient compared to wildtype cells (Fig. 4*B*), recapitulating the findings described above with EPO-eGFP and ruling out an indirect effect due to the eGFP tag.

**Fig. 4.**
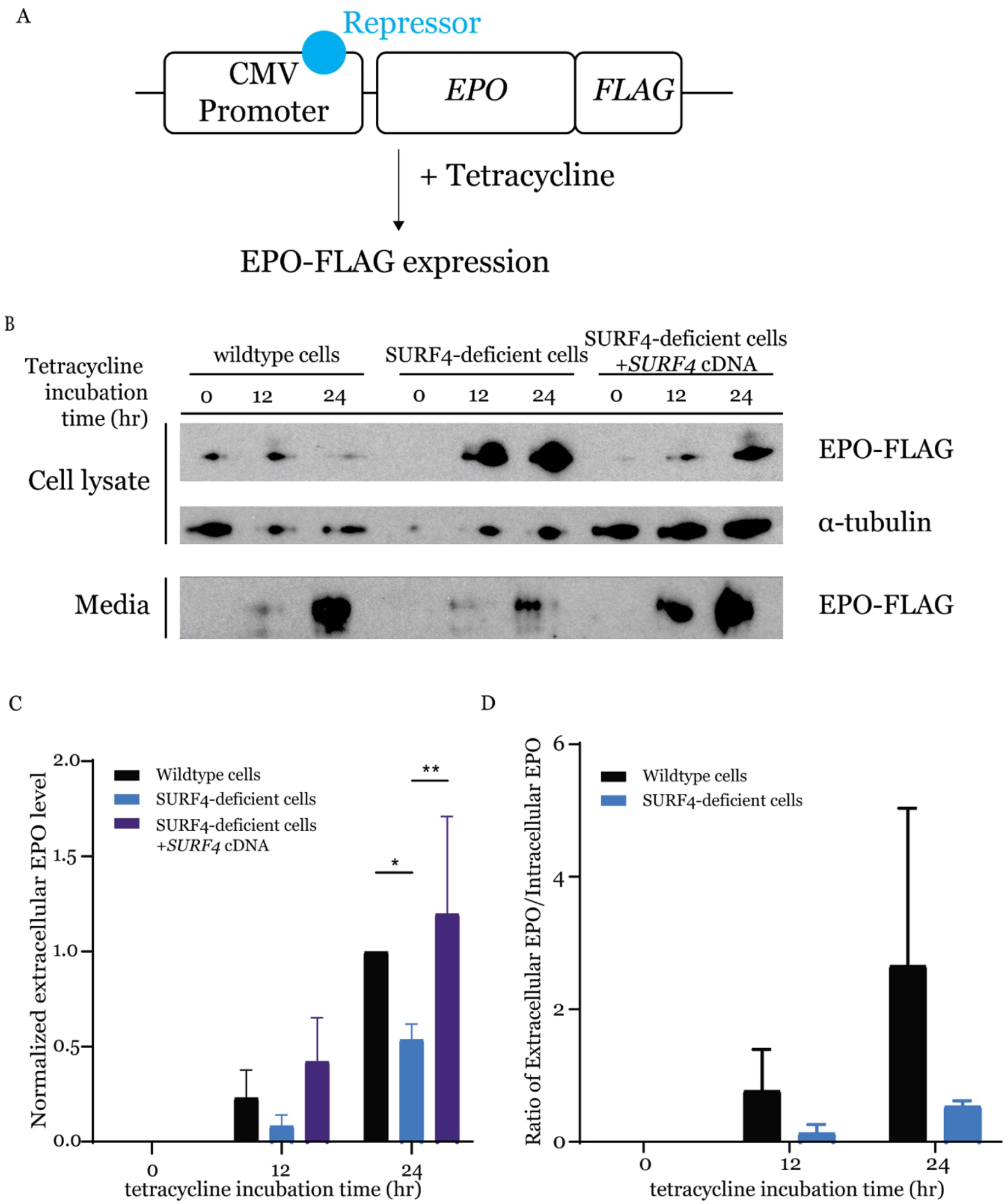
*SURF4* mutagenesis causes reduced Extracellular EPO-FLAG secretion. (*A*) We generated a FLP-In TREX HEK293 cell line with tetracycline inducible EPO-FLAG expression. (*B*) Intracellular and extracellular EPO-FLAG abundance in wild-type, SURF4-deficient, and SURF4-rescued cells was measured by Western blot (using anti-FLAG antibody) after 0, 12, and 24 hours of incubation with tetracycline. α-tubulin was used as loading control. (*C*) Quantification of densitometry of extracellular EPO and (*D*) ratios of extracellular/intracellular EPO normalized to α-tubulin in 3 independent experiments. * p<0.05, ** p<0.01 by two-way ANOVA. Error bars represent standard deviations.

SURF4 localizes to the ER membrane (31–33) and functions as an ER cargo receptor, suggesting that the increased accumulation of intracellular EPO in the setting of SURF4 deficiency is secondary to reduced EPO secretion. Consistent with this hypothesis, the extracellular EPO-FLAG protein level was considerably lower in the conditioned media of SURF4-deleted cells compared to wildtype cells (Fig. 4 *B* and *C*), as was the ratio of extracellular to intracellular EPO-FLAG levels (Fig. 4*D*). The latter findings observed in SURF4-deficient cells were rescued by stable expression of *SURF4* cDNA (Fig. 4 *B* and *C*). These results indicate that disruption of *SURF4* results in a defect in EPO secretion.

### *SURF4* deletion results in accumulation of EPO in the ER

We next performed live cell fluorescent confocal microscopy to determine the localization of accumulated EPO in the setting of *SURF4* deletion. We co-transfected the EPO-eGFP/A1AT-mCherry reporter construct (Fig. 1*A*) with a vector expressing an ER blue fluorescent marker (ERoxBFP) into wildtype or SURF4 deficient (*SI Appendix*, Fig. S1) HEK293 cells. We quantified the degree of co-localization between EPO and ERoxBFP (as well as A1AT and ERoxBFP, as control) by Pearson correlation coefficient (PCC). SURF4 deficient cells exhibited an increased co-localization of EPO (but not A1AT) with ERoxBFP compared to wildtype cells (PCC 0.7870 in *SURF4* deleted cells versus 0.2934 in wildtype cells, p<0.0001) (Fig. 5 *A* and *B*).

**Fig. 5.**
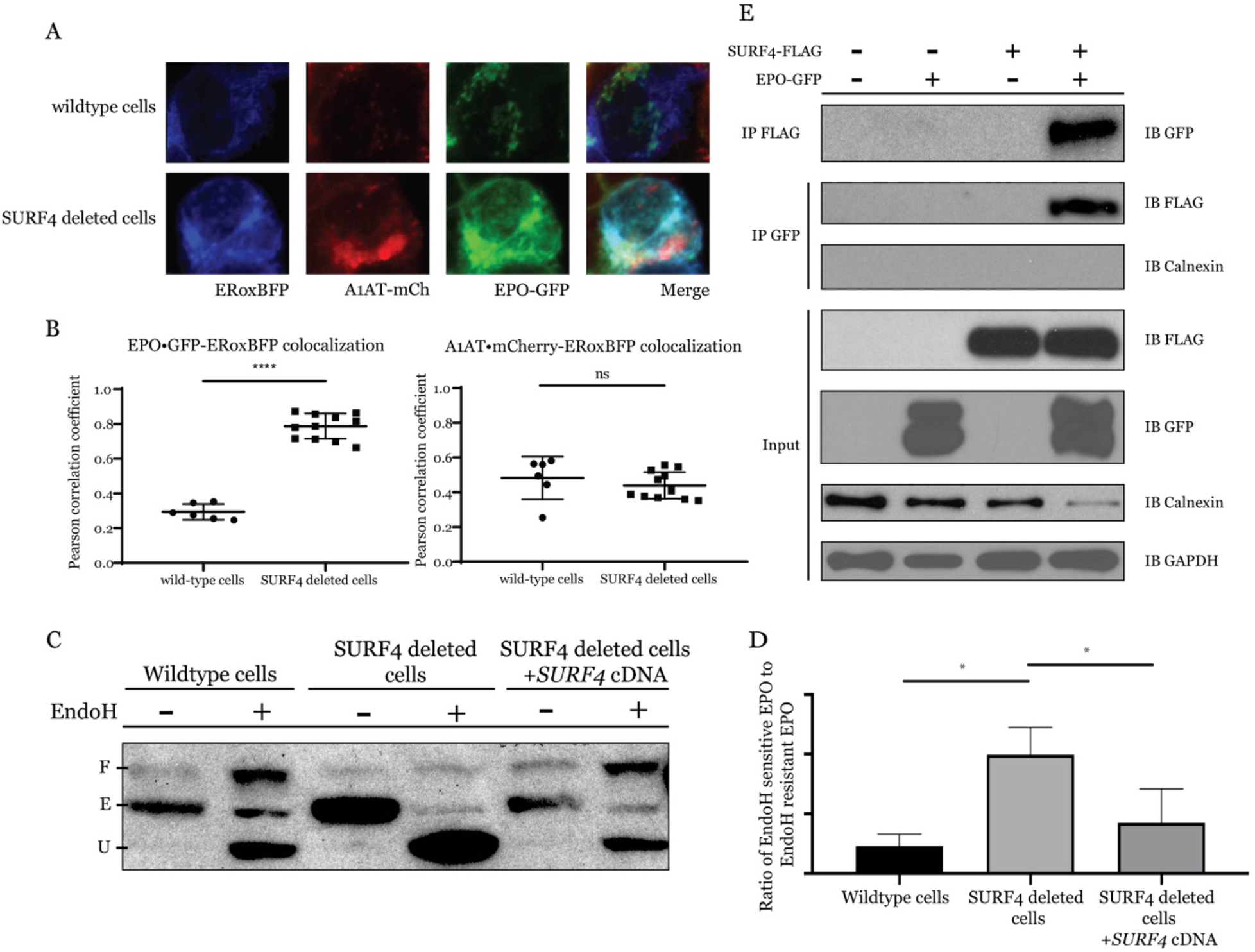
Disruption of *SURF4* results in accumulation of EPO in the ER. (*A*) Live cell fluorescent confocal microscopy of wildtype or *SURF4* deleted reporter cells expressing the ER marker, ERoxBFP. (*B*) Quantification of the degree of co-localization between EPO and ERoxBFP, as well as A1AT and ERoxBFP as control, by Pearson correlation coefficient. n=6 for wildtype, n=12 for SURF4 deficient cells. **** p-value <0.0001, unpaired student t-test, ns = non-significant. (*C*) Cell lysates were collected from wildtype, *SURF4* deleted, or *SURF4* rescued cells (*SURF4* deleted cells with stable expression of wildtype *SURF4* cDNA) expressing EPO-eGFP and were either treated with EndoH or left untreated. Immunoblotting was done with anti-GFP antibody. E = ER form of EPO (endoH sensitive), U = unglycosylated EPO, F = fully glycosylated EPO (post-Golgi form of EPO) as demonstrated by treating wildtype cells with either PNGase or EndoH (see Fig. S2). (*D*) Quantification of EndoH sensitivity from 3 independent experiments. * p<0.05. (*E*) FLAG antibody or eGFP antibody were used to immunoprecipitate EPO-eGFP or SURF4-FLAG, respectively, from lysates of cells expressing either EPO-eGFP, SURF4-FLAG, both, or neither.

To confirm the ER accumulation of EPO upon *SURF4* disruption, we tested the glycosylation status of EPO in SURF4-deficient cells. EPO contains 3 N-glycosylation sites. In the ER, N-linked high mannose oligosaccharides are added to EPO and further modifications are made in the Golgi apparatus. The ER form of EPO is cleavable by EndoH, but the post-ER form is not. Therefore, the ratio of EndoH cleaved to EndoH uncleaved EPO will approximate the ratio of the amount of EPO in the ER versus the amount of EPO in the Golgi apparatus or beyond. In SURF4 deficient cells, the ratio of ER/post-ER form of EPO was significantly increased compared to that in wildtype cells (Fig. 5 *C* and *D* and *SI Appendix*, Fig. S2), an effect that was decreased by stable expression of *SURF4* cDNA (Fig. 5 *C* and *D*). Taken together, these results demonstrate that SURF4 promotes the efficient ER exit and secretion of EPO.

### SURF4 physically interacts with EPO

To determine if SURF4 binds to EPO, we tested for reciprocal co-immunoprecipitation of SURF4-FLAG and EPO-GFP in HEK293T cells. An antibody against the FLAG epitope co-immunoprecipitated EPO-eGFP but not the ER luminal resident protein calnexin. Similarly, an antibody against GFP co-immunoprecipitated FLAG-SURF4 (Fig. 5*E*). These results are consistent with a specific physical interaction between SURF4 and EPO.

TPO shares significant sequence homology with EPO. To test if TPO, similarly to EPO, depends on SURF4 for efficient secretion, we generated 2 independent clonal HEK293 cells stably expressing and efficiently secreting TPO-eGFP and A1AT-mCherry (Fig. 6 *A* and *B*). Like A1AT, TPO did not accumulate intracellularly upon *SURF4* deletion (Fig. 6 *C* and *D*). These findings demonstrate the specificity of SURF4 for promoting EPO secretion and suggest that the SURF4/EPO interaction is mediated by one of the EPO domains not present in TPO.

**Fig. 6.**
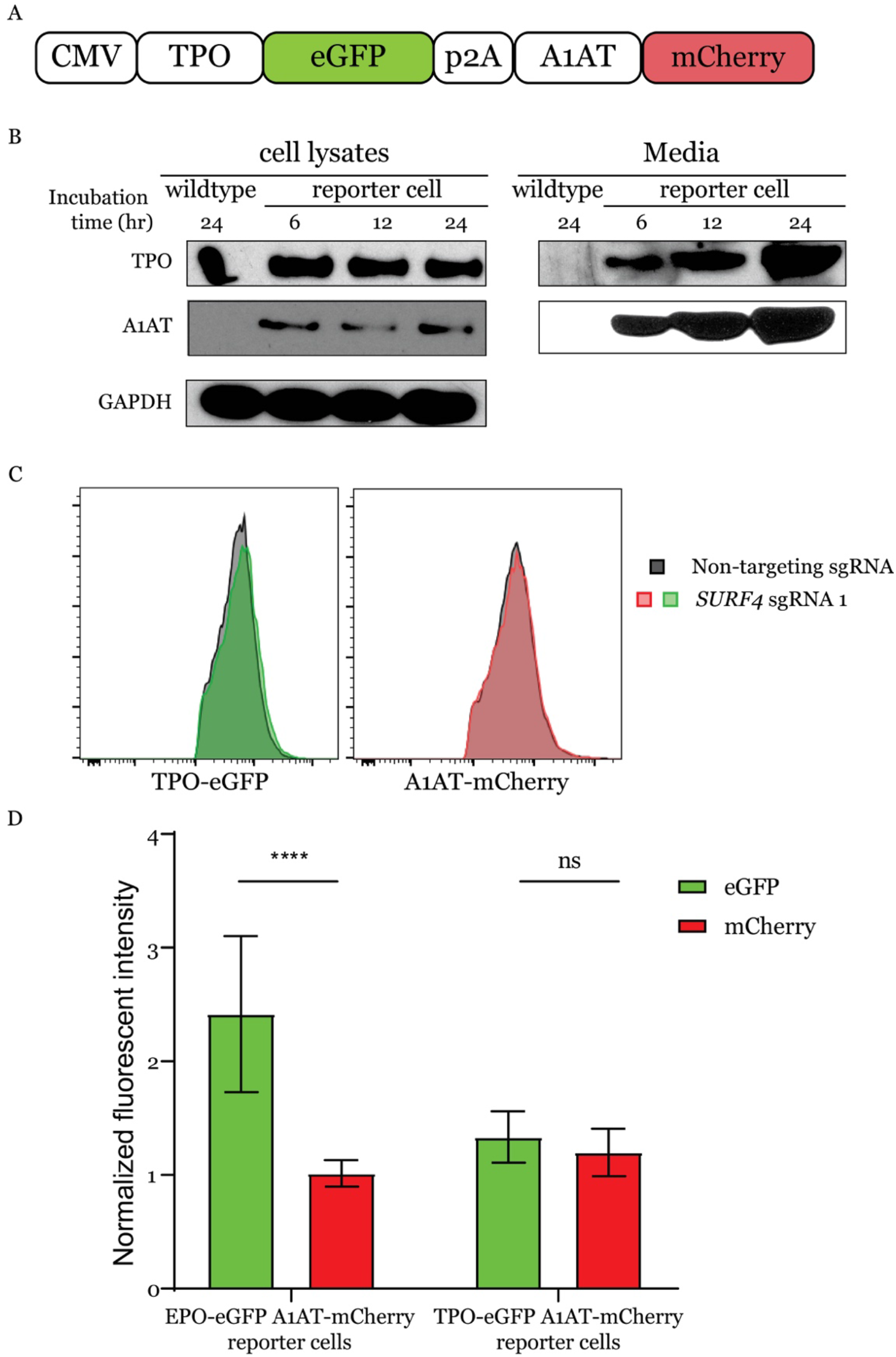
Thrombopoietin secretion does not depend on SURF4. (*A*) A construct that expresses TPO-eGFP and A1AT-mCherry from the same CMV promoter was assembled and used to generate a reporter cell line stably expressing these two fusion proteins. (*B*) Intracellular and extracellular TPO-eGFP and A1AT-mCherry protein abundance was determined by Western blot using anti-eGFP and anti-mCherry antibodies, respectively. (*C*) Flow cytometry histograms showing absence of intracellular accumulation of TPO following *SURF4* deletion in HEK293T cells. (*D*) Quantification of cellular mean flourescence intensity of TPO-eGFP and A1AT-mCherry in cells transduced with *SURF4*-targeting sgRNAs (n=29). Results were normalized to mean flourescence intensity of cells transduced with non-targeting sgRNAs. As a positive control, the same experiment was performed in parallel in reporter cell lines expressing EPO-eGFP and A1AT-mCherry (n=48). **** p<0.0001, ns=non-significant.

### *SURF4* promotes the secretion of endogenous EPO

The experiments described above were performed in a heterologous cell line overexpressing EPO fused to either an eGFP or a FLAG tag. To test the impact of *SURF4* deletion on the secretion of endogenous EPO, we transduced human HEP3B cells with *SURF4*-targeting sgRNAs or control sgRNAs. As a positive control, a sgRNA targeting *EPO* resulted in profound reduction of extracellular EPO level to almost an undetectable (0.45% of control) level (Fig. 7). Disruption of *SURF4* in HEP3B cells using 2 independent sgRNAs resulted in a significant reduction (51.22% of control) of extracellular EPO levels compared to cells transduced with control sgRNAs (Fig 7).

**Fig. 7.**
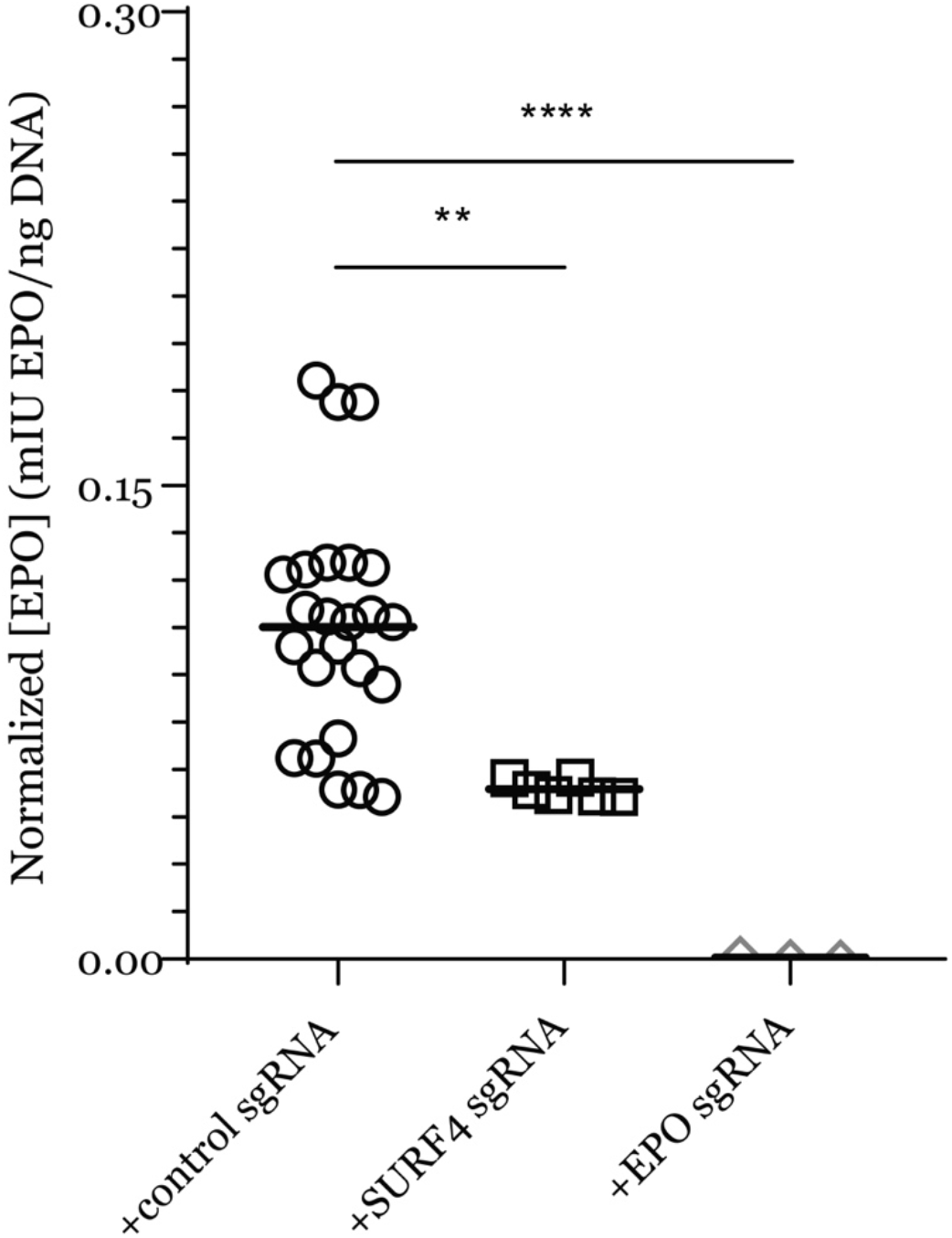
*SURF4* deletion in HEP3B cells results in reduced extracellular secretion of EPO expressed from its endogenous genomic locus. HEP3B cells were transduced with lentivirus expressing *SURF4*-targeting sgRNAs, control sgRNAs, or EPO-targeting sgRNA as a positive control. EPO expression from its endogenous regulatory elements was subsequently induced with CoCl_2_ and measured in the conditioned media by ELISA and normalized to the total number of cells. ** p <0.01, **** p<0.0001.

### SURF4 overexpression promotes more efficient EPO secretion

We next determined if SURF4 overexpression promotes more efficient EPO secretion. We generated a lentivirus expressing equal amounts of SURF4 and Katushka2S (SURF4-p2A-Katushka2S, Fig. 8*A*) and transduced it into HEK293 cells expressing EPO-FLAG from a tetracycline inducible promoter. Cells with the highest (top 10%) and lowest (bottom 10%) SURF4 expression, as determined by Katushka2 fluorescence, were FACS sorted. Following tetracycline administration, EPO level was found to be significantly increased in the conditioned media of cells overexpressing SURF4 compared to cells expressing low SURF4, with the reverse pattern observed intracellularly (Fig. 8 *B*-*D*).

**Fig. 8.**
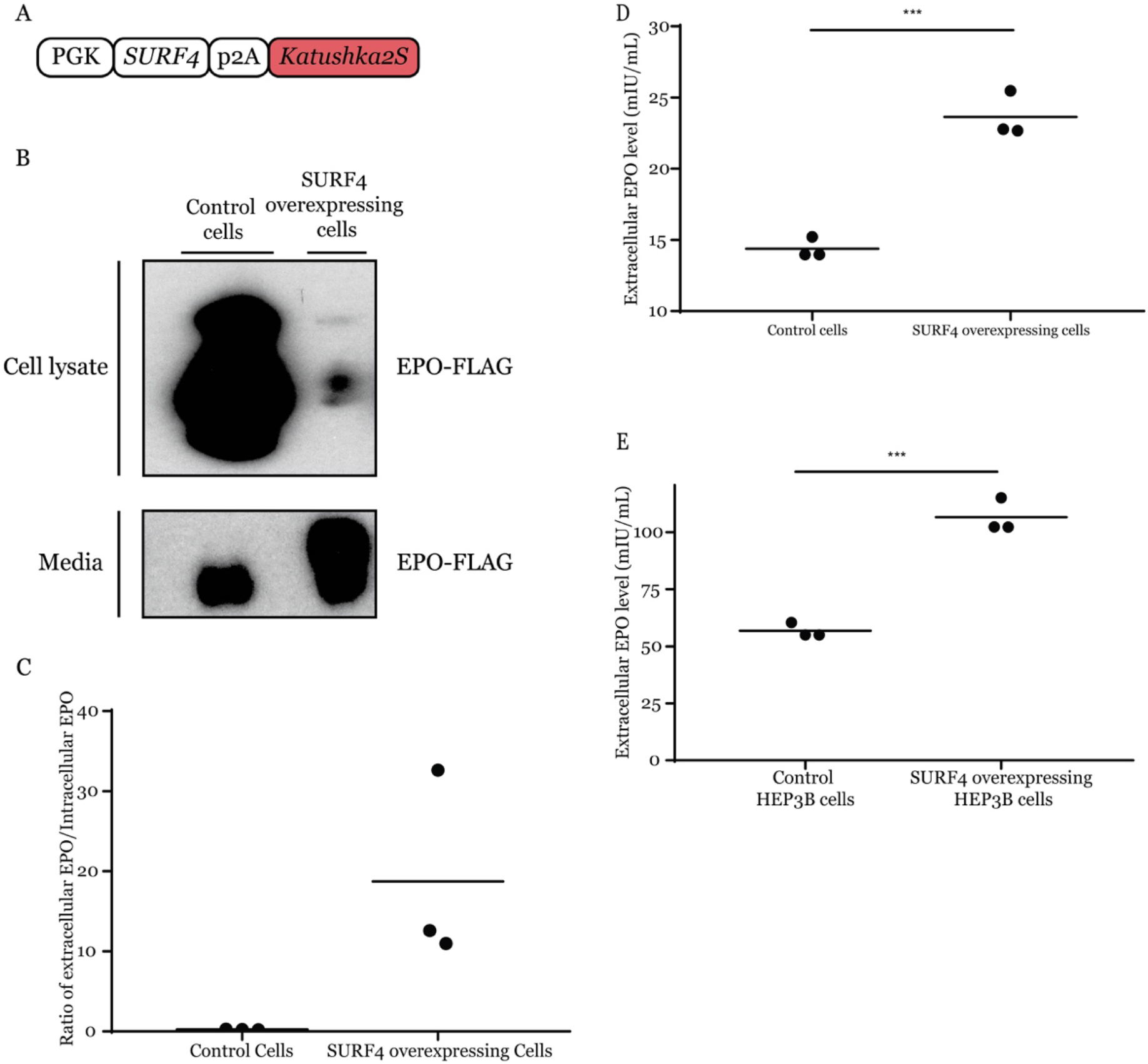
SURF4 overexpression leads to enhanced EPO secretion. (*A*) A lentiviral construct that expresses equal amounts of SURF4 and Katushka2S from the same PGK promoter was assembled and transduced into HEK293 cells expressing EPO-FLAG from a tetracycline inducible promoter. Cells with top 10% and bottom 10% Katushka2S fluorescence were FACS sorted, corresponding to cells overexpressing SURF4 and control cells, respectively. (*B*) Intracellular and extracellular EPO abundance following a 12-hour tetracycline incubation was analyzed by Western blot (using anti-FLAG antibody) and (*C*) quantification of densitometry of the ratio of extracellular/intracellular EPO was determined in 3 independent experiments. (*D*) The extracellular EPO level was also measured by ELISA. (*E*) HEP3B cells overexpressing SURF4 (and control cells) were generated as described above. Following incubation with CoCl_2_, the extracellular EPO level was measured by ELISA. ***P<0.001, unpaired t-test.

To assess the impact of SURF4 overexpression on the secretion of EPO expressed from its endogenous genomic locus, we performed the same experiment described above in HEP3B cells. EPO level was increased in the conditioned media of cells expressing high compared to low SURF4 levels (Fig. 8*E*). Taken together, these results demonstrate that SURF4 overexpression promotes more efficient EPO secretion.

## Discussion

In this report, we developed an unbiased genome-scale loss-of-function screen and identified *SURF4* as a regulator of intracellular EPO levels. Deletion of *SURF4* resulted in intracellular accumulation and extracellular depletion of EPO. Overexpression of SURF4 resulted in the reversed pattern. Consistent with the reported localization of SURF4 at the ER membrane (32, 34, 35), we found that disruption of *SURF4* resulted in accumulation of EPO in the ER, and that EPO and SURF4 physically interact. Taken together, these results strongly suggest that SURF4 is the ER cargo receptor that mediates the efficient secretion of EPO.

The screen described above was performed in a cell line with heterologous overexpression of EPO under the control of a CMV promoter. Therefore, it was important to examine if SURF4 facilitates the secretion of EPO when expressed at a more physiological level. Accordingly, we deleted *SURF4* in HEP3B cells which were induced to express EPO from its endogenous genomic locus, and found that SURF4 also promotes EPO secretion under these conditions.

SURF4 is the mammalian ortholog of yeast Erv29p. Erv29p facilitates packaging of pro-alpha-factor in COPII vesicles promoting their ER-to-Golgi transport (31, 36, 37). Erv29p recycles back from Golgi-to-ER via recognition of its well-conserved di-lysine sorting signal by the COPI coat (38). In mammalian cells, only a handful of cargos (APOB, PCSK9, DSSP, AMLEX, and GH) have been shown to depend on SURF4 for efficient secretion (32, 33, 39). However, a recent report demonstrated that mice with germline deletion of *Surf4* exhibit early embryonic lethality (40) similar to *C. elegans* (33), suggesting the presence of one or more SURF4-dependent cargos with a critical function during embryogenesis. Future studies aimed at identifying the repertoire of cargos that depend on SURF4 for secretion are essential.

A recently published report suggested that the cargo proteins that depend on SURF4 for secretion contain an N-terminal tripeptide ‘ER-ESCAPE’ motif (39). This motif is exposed following removal of the leader sequences and is recognized by SURF4 (39). However, an N-terminal ‘ER-ESCAPE’ motif with high SURF4 binding affinity is not present in EPO. Additionally, we found that EPO, but not TPO, depends on SURF4 for efficient secretion. However, TPO has an N-terminal motif with a better predicted SURF4 binding affinity than EPO. These results suggest that the N-terminal ‘ER-ESCAPE’ motif may not be a generalizable determinant of SURF4 interaction for all SURF4-dependent cargos.

Soluble cargos are exported from the ER via the passive “bulk flow” or the concentrative “cargo capture” processes, with several lines of evidence supporting one route versus the other(4). Though “bulk flow” and “cargo capture” are not mutually exclusive, this report provides support for the “cargo capture” model of EPO secretion. However, it is important to note that in our experimental conditions, ~50-70% of extracellular EPO is reduced in the setting of SURF4-deficiency. Therefore, the secretion of the remaining EPO depends on either bulk flow or one or more separate and unidentified receptors.

Recent developments in genome engineering using CRISPR/Cas9 technology have dramatically enhanced the potential and efficacy of comprehensive, high throughput genetic screens (41–55). Such strategies can be applied *in vitro* and *in vivo* to discover novel biologic insights. Our screen was designed to focus on post-transcriptional regulators of EPO by placing its expression under the control of a CMV promoter. Screening strategies similar to the one employed in this manuscript and in a recently published report (32) might help identify additional ER cargo receptors for other soluble secreted proteins, and shed more light into the extent of the contribution of “cargo capture” to recruitment of cargos into COPII vesicles.

Findings in this report may have important implications for erythropoiesis. EPO, the master regulator of erythropoiesis, is produced by specialized peritubular fibroblasts in the kidney. The transcriptional control of EPO via the hypoxia inducible factor pathway has been well studied (28, 56–65) culminating in the development of prolyl hydroxylase inhibitors, a class of compounds that increase EPO production at the transcriptional level via activation of the hypoxia inducible factor (66–73). These drugs are currently in clinical development, with several compounds in advanced phase 2 or 3 trials (74–78); however, there are numerous potential concerns and adverse effects of these drugs, including possible increased risks of malignancy and autoimmune disease (79–81). Similar to the transcriptional control of EPO, the intracellular signal transduction pathway downstream of the EPO receptor has also been well studied(82–84). In contrast, much less is known about the molecular basis of EPO trafficking. Our findings suggest that modulating SURF4 may be effective for the treatment of disorders of erythropoeisis that are driven by aberrant EPO levels (85–90). Though a handful of other cargos depend on SURF4 for their secretion (32, 33, 39), with additional cargos likely remaining to be identified, targeting SURF4 exclusively in the EPO producing cells might alter plasma EPO levels and therefore regulate erythropoiesis without affecting other SURF4-dependent cargos that are expressed in other cells. Alternatively, an inhibitor that specifically disrupts the SURF4-EPO interaction would also be expected to have no effects on other cargos that bind SURF4.

Recombinant human EPO (rhEPO) is used clinically for the treatment of anemia due to chronic kidney disease, chemotherapy, or ziduvidine. rhEPO is also used to reduce the requirement of allogeneic red blood cell transfusion following certain elective surgeries. Though the use of rhEPO is indicated in only a subset of the above clinical scenarios, the rhEPO market size was valued at ~7.4 billion US dollars in 2016 (91). In this report, we demonstrate that SURF4 overexpression results in enhanced EPO secretion. This approach could be applied to increase the efficiency of rhEPO production, which might translate into reduced costs of this drug.

## METHODS

### Cell culture

HEK293T and HEP3B cells were purchased from ATCC. Flp-In T-REx 293 cells were purchased from Invitrogen. HEK293T and Flp-In T-REx 293 cells were cultured in DMEM (Gibco) supplemented with 10% heat-inactivated fetal bovine serum (Peak) and 1% penicillin-streptomycin (Gibco). HEP3B cells were cultured in alpha-MEM (Gibco) supplemented with 2mM L-glutamine (Gibco), 10% heat-inactivated fetal bovine serum (Peak), and 1% penicillin-streptomycin (Gibco). All cells were grown in a humidified 37°C incubator with 5% CO_2_.

### Generation of the EPO-GFP A1AT-mCherry reporter cell line

The CMV-EPO-eGFP-p2A-A1AT-mCherry construct was assembled using the NEBuilder HiFi DNA assembly cloning kit (NEB) using vector sequences derived from PCSK9-eGFP-p2A-A1AT-mCherry (32) and EPO cDNA obtained from Dharmacon. HEK293T cells were transfected with CMV-EPO-eGFP-p2A-A1AT-mCherry using Fugene HD transfection reagent (Promega). Transfected cells were selected with 350µg/mL hygromycin (Invitrogen). Five weeks following hygromycin selection, single cells were sorted into 96-well plates using a SY-3200 flow cytometer (Sony). Single cell clones were expanded and analyzed for stable expression of EPO-eGFP and A1AT-mCherry using a LSR Fortessa flow cytometer (BD Bioscience).

### Generation of the TPO-GFP A1AT-mCherry reporter cell line

The TPO cDNA sequence was amplified from human liver RNA (de-identified tissue sample obtained from the tissue procurement core, University of Michigan, IRB #HUM00048303) and the CMV-TPO-eGFP-p2A-A1AT-mCherry construct was generated and transfected into HEK293T cells as described in the paragraph above. Single cell clones were sorted, expanded, and analyzed for stable expression of TPO-eGFP and A1AT-mCherry, as described above.

### Expansion and lentiviral preparation of the pLentiCRISPRv2 library

The pLentiCRISPRv2 whole genome CRISPR library was obtained from Addgene (Addgene #1000000048, a gift from Feng Zhang (41)), expanded by 16 electroporations (8 for each half library) into Endura electrocompetent cells (Lucigen), and plated on sixteen 24.5 cm bioassay plates (ThermoFisher Scientific). Following a 12-14 hour incubation at 37C, colonies were harvested from agar plates, and pooled plasmids for each half library were isolated separately by Maxipreps using an EndoFree Plasmid Maxi Kit (Qiagen). To prepare the pooled lentiviral library, 11.3 ug of each half library was co-transfected with 17 ug of psPAX2 (Addgene #12260, a gift from Didier Trono) and 11.3 ug of pCMV-VSV-G (addgene #8454, a gift from Robert Weinberg (92)) using Lipofectamine LTX with PLUS reagent (ThermoFisher Scientific) into each of six T225 tissue culture flasks (ThermoFisher Scientific) containing HEK293T cells at ~80-90% confluency. Media was changed 24 hours post-transfection, and viral supernatant was collected 12, 24, and 36 hours afterwards. Media containing viral supernatant was centrifuged at 500 g for 5 min, pooled, aliquoted, snap-frozen in liquid nitrogen, and stored at −80C.

### CRISPR/Cas9 loss-of-function genome wide screen

For each independent screen, more than 142 million reporter cells were plated in 15-cm tissue-culture dishes (Corning) at 30% confluency. Cells were transduced with the lentiviral library (with 8ug/ml polybrene, Sigma) at a multiplicity of infection (MOI) of ~0.3. Twenty-four hours post-viral transduction, puromycin selection (1 ug/ml, Sigma) was applied for 4 days. Subsequently, cells were kept at a logarithmic phase of growth and passaged every 2-3 days, maintaining more than 36 million cells in culture at all times in order to preserve library depth. Fourteen-days post-transduction, ~80 million cells were isolated from tissue culture dishes using trypsin 0.25% (Gibco), pelleted by centrifugation (350g, 4C, 5 min), resuspended in cold PBS + 2% FBS, and filtered through a 35 um mesh into flow cytometry tubes (Corning). Cells were divided into 20 tubes and maintained on ice until sorting. Cells with normal mCherry fluorescence (mid 70-80% fluorescence) and top or bottom ~7% eGFP fluorescence (~4 million cells/population) were sorted using a BD FACSAria III (BD Biosciences) and collected into 15 ml polypropylene tubes (Cellstar) containing media. Genomic DNA was extracted using a DNeasy Blood & Tissue kit (Qiagen), and integrated lentiviral sgRNA sequences were amplified by a two-step PCR reaction (20 cycles and 14 cycles, respectively) as previously described (32, 41) using a Herculase II Fusion DNA Polymerase kit (Agilent biotechnologies). DNA was purified after each of the PCR reactions using a QIAquick PCR purification kit (Qiagen). Following the 2 step PCR, DNA was analyzed with a bioanalyzer (Agilent) and the sgRNA amplicons were sequenced using a NextSeq 500 Sequencing System (Illumina). On average, 23.5 million reads were generated for each sorted cell population of each screen. Overall, 98% of the reads had a per sequence quality score (phred-based base quality score) of greater than 30. 104,331 sgRNA sequences were mapped and identified (along with the barcode corresponding to each cell population of each replicate) using a custom Perl Script as previously described (32). Enrichment at the sgRNA and gene levels was analyzed using DESeq2 and MAGeCK, respectively (93, 94).

### Disruption of candidate genes using CRISPR/Cas9

sgRNAs targeting several genes and several non-targeting sgRNAs (listed in *SI Appendix*, Table S1) were cloned into the pLentiCRISPRv2 plasmid (Addgene:52961, a gift from Feng Zhang (41) as previously described (42). pLentiCRISPR plasmids were packaged into Lentivirus, using the same method described above. To disrupt genes in a population of cells, cells were transduced with lentivirus at an MOI of ~0.3. Subsequently, transduced cells were selected with puromycin and passaged for 10-14 days prior to FACS analysis. For all validation experiments, a minimum of 3 biologic replicates were analyzed.

### Generation of SURF4-deficient clonal cell lines

To generate clonal cell lines that are deficient for SURF4, a sgRNA targeting *SURF4* exon 2 (*SI Appendix*, Table S1) was cloned into the PX459 plasmid (Addgene: 62988, a gift from Feng Zhang) as previously described (95), and the construct was transiently transfected into cells using Fugene HD transfection reagent (Promega). Twenty-four hours post-transfection, puromycin (1ug/mL, Sigma) selection was applied for 3 days, and subsequently, single cells were sorted into each well of three 96-well plates using the SY-3200 flow cytometry instrument (Sony). Following expansion of the single cell clones, genomic DNA was extracted with QuickExtract (Epicentre) and indels were determined by amplification of the sgRNA target site by polymerase chain reaction using Herculase II Fusion DNA Polymerase (Agilent biotechnologies) and Sanger sequencing. Primers used for PCR and Sanger sequencing are listed in *SI Appendix*, Table S1. Three independent single cell clones with homozygous frameshift indels in *SURF4* were generated.

### Flow Cytometry analysis

HEK293T cells were detached with 0.25% trypsin (Gibco), washed with PBS, collected by centrifugation (350 g, 5 min, 4C), resuspended in cold PBS with 0.1% BSA and 10mM HEPES (Invitrogen), filtered with 70 μm cell strainers, and analyzed by BD LSR Fortessa (BD Bioscience). FlowJo (Tree Star) was used to calculate the mean fluorescence intensity and for further analysis.

### Brefeldin A treatment

HEK293 cells stably expressing EPO-eGFP and A1At-mCherry were incubated with 1ug/mL Brefeldin A (Biolegend) for 12 hr. Subsequently, cells were collected as described above and analyzed by flow cytometry for intracellular accumulation of EPO-eGFP and A1AT-mCherry.

### Western blots

To prepare cell lysates, cells were washed in PBS, suspended in RIPA buffer (Invitrogen) supplemented with cOmplete protease inhibitor cocktail (Sigma), briefly sonicated, and incubated for 30 minutes in the cold room with end-over-end rotation. Cell lysates were cleared by centrifugation to remove cell debris (20,000 g, 30 min, 4C) and were analyzed immediately or stored at −80C until analysis. Protein quantification was performed using Pierce BCA protein assay kit (ThermoFisher Scientific) per manufacturer’s instruction. Lysates were boiled for 5 min at 95C with 4X Laemmli sample buffer (Bio-Rad) supplemented with β-mercaptoethanol. Equal amounts of proteins were loaded on either a 4-12% Bis-Tris gel or a 4-20% Tris-Glycine gel (Invitrogen), and SDS gel electrophoresis was performed as previously described (96, 97). Proteins were then transferred onto a nitrocellulose membrane (Bio-Rad). Following blocking in 5% (wt/vol) milk-Tris-buffered saline with Tween (TBST), membranes were incubated with primary antibody at 4C overnight, washed 3 times in TBST, probed with peroxidase-coupled secondary antibodies, washed again 3 times in TBST, and developed with SuperSignal West Pico Plus (ThermoFisher Scientific). For HRP-conjugated primary antibodies, nitrocellulose membranes were incubated with these antibodies and immediately developed following 3 TBST washes. Densitometry was performed with ImageJ. To test for the secretion efficiency of various cargo proteins, cells were seeded at equal densities in 6-well plates or 10-cm plates, and conditioned media was collected at different time points, cleared by centrifugation (500 g, 5 min, 4C), and analyzed immediately (by western blot) as described above, or stored at −80C until analysis.

### Antibodies

The following antibodies were used for immunoblotting: Anti-GFP (Abcam, ab290), anti-mCherry (Abcam, ab167453), anti-Calnexin (Cell Signaling, 2679S), anti-GAPDH (Millipore, MAB374), horseradish peroxidase (HRP) conjugated anti-FLAG (Abcam, ab1238), anti-alpha-tubulin (Abcam, ab176560), HRP conjugated anti-mouse IgG (Biorad, 1706516) and HRP conjugated anti-rabbit IgG (Jackson ImmunoResearch Laboratories, 111-035-003).

### Tetracycline induced EPO-FLAG expression

The coding sequence of EPO with a C-terminal Flag was cloned into pDEST-pcDNA5-BirA-FLAG (Invitrogen) using NEBuilder HiFi DNA assembly cloning kit (NEB). Wildtype, SURF4-deficient (with homozygous frameshift *SURF4* indels), or SURF4-rescue (with homozygous frameshift *SURF4* indels but with stable expression of *SURF4* cDNA) Flp-In T-REx HEK293 cells with tetracycline-inducible expression of EPO-FLAG were generated as previously described (98). To induce the expression of EPO-FLAG, tetracycline (1 ug/mL) was added to the media. Cells and media were collected prior to the addition of tetracycline and 12- and 24-hours following tetracycline. Intracellular and extra-cellular EPO levels were analyzed by western blot as described above.

### Endoglycosidase H (EndoH) assay

HEK293T cells that are either wild-type, SURF4-deficient, or SURF4-rescue (defined in the paragraph above) were transfected with a plasmid expressing EPO-eGFP. Thirty-six hours post-transfection, total cell lysates were prepared, and protein quantification was performed, both as described above. Lysates were incubated with denaturing buffer (NEB) 95°C for 10 minutes and equal amounts of lysates (180 ug) were treated with either 1uL of EndoH (NEB), PNGase F (NEB), or DMSO as control for 1 hour (37°C). Subsequently, Laemmli buffer (Bio-RaD) was added and the samples were boiled (95°C) for 5 min. Samples were loaded on a 4-12% Bis-Tris Gel (Invitrogen) and Western blot was performed as described above. This experiment was performed in biologic triplicates.

### Live cell confocal fluorescent microscopy

Wild-type or SURF4-deficient HEK293T cells that stably express EPO-eGFP and A1AT-mCherry were transfected with a plasmid expressing ERoxBFP (Addgene: 68126, a gift from Erik Snapp (99)). Twenty-four hours post-transfection, cells were seeded on Lab-Tek Chambered Coverglass (ThermoFisher). Fluorescent images were captured on a Nikon A2 confocal microscope. To quantify colocalization between 2 proteins, Pearson correlation coefficient was calculated using the Nikon Elements software. This experiment was performed and analyzed by an investigator blinded to the genotype of the cells.

### Co-immunoprecipitation

Flp-In T-Rex 293 cells that are either wild-type, SURF4-deficient, or SURF4-rescue (defined above) were transfected with CMV-EPO-eGFP-p2A-A1AT-mCherry using Fugene HD transfection reagent (Promega). Twenty-four hours post-transfection, cells were washed with PBS and incubated in PBS containing 2 mM dithiobis (succinimidyl propionate) (Pierce) for 30 min at room temperature. Subsequently, 20 mM Tris-HCL (pH 7.5) was added to quench the reaction. Cells were then washed twice in PBS and cell lysis was performed with the following lysis buffer (100 mM tris, 10% glycerol, 1% NP-40, 130 mM NaCl, 5 mM MgCl_2_, 1 mM NaF and 1mM EDTA, supplemented with cOmplete protease inhibitor cocktail pH 7.5). Cell lysates were collected as described above and incubated overnight at 4C with either anti-FLAG M2 magnetic beads (Sigma) or GFP-Trap beads (ChromoTek). Following 5 washes with lysis buffer, proteins were eluted from the beads via incubation with 2X Laemmli sample buffer containing β-mercaptoethanol for 15 minutes at room temperature.

### Generation of cell lines expressing low or high SURF4 levels

A construct expressing SURF4 and the Katushka2S fluorescent marker (PGK-SURF4-p2A-Katushka2S) was assembled with the NEBuilder HiFi DNA assembly cloning kit (NEB) using vector sequence derived from LV1-5 (Addgene #68411) and cDNAs of *SURF4* and Katushka2S (a gift from Gary Luker(100)). The construct was packaged into lentivirus as described above and transduced at MOI of ~1 into Flp-In T-REx 293 or HEP3B cells. Transduced cells were selected with puromycin and passaged for 14 days prior to FACS sorting. Cells with top and bottom 10% Katushka2S fluorescence we sorted.

### Generation of SURF4-deficient HEP3B cells

Wild-type HEP3B cells were transduced with lentiviral sgRNA targeting *SURF4*, control sgRNA (combination of non-targeting sgRNAs and sgRNAs targeting genes that do not affect EPO: *BCL11A*, *MPL*, *SERPINA1*), or sgRNA targeting *EPO* as a positive control. Cells were selected with puromycin and passaged for at least two weeks prior to further analysis. EPO levels in the conditioned media were compared between *SURF4* deleted cells and control cells, correcting for the total cell number at the time of EPO measurement. Genomic DNA was extracted from HEP3B cells using QuickExtract (Epicentre).

### EPO ELISA

Equal numbers of cells were seeded in 6-well or 24-well plates. For HEP3B cells, EPO production was stimulated with CoCl_2_ (75 μM, Sigma) for 24 hours and conditioned media was collected and cleared by centrifugation (500 g, 5 min, 4C). For Flp-In T-Rex HEK293 cells with tetracycline-inducible expression of EPO-FLAG, tetracycline (1µg/mL) was added for 12 hours and conditioned media was collected, cleared by centrifugation (500 g, 5 min, 4C) and diluted 1:500. EPO level was measured in the conditioned media using the LEGEND MAX Human Erythropoietin ELISA kit (Biolegend), according to manufacturer’s instructions.

### Statistical analysis

CRISPR screen data analysis was performed as described above. The statistical differences in mean fluorescence intensity between EPO-eGFP and A1AT-mCherry were compared by two-way ANOVA. The difference in extracellular EPO-FLAG level amongst wild-type, SURF4-deficient, and SURF4-rescued Flp-In T-REx 293 cells were compared by two-way ANOVA. The Pearson correlation coefficient differences between wildtype and SURF4 deficient HEK293T cells were compared by unpaired t-test. The statistical difference in extracellular EPO detected by EPO ELISA was assessed using an unpaired t-test. The difference in relative amount of EndoH sensitive EPO amongst wildtype, SURF4-deficient, and SURF4-rescued HEK 293T cells was assessed by one-way ANOVA.

## Acknowledgments

This work was supported by National Institute of Health Grants K08 HL128794 (R.Kh.) and R01 HL148333 (R.Kh.). This work was also supported by MCubed, a research seed-funding program for faculty at the University of Michigan (R.Kh., K.D.), and by the University of Michigan Rogel Cancer Center (R.Kh.). R.Ki. was supported by NIH T32-CA009357. G.B.C. was spported by NIH T32-GM007315.

## Author Contributions

Z.L. and R.Kh. conceived the study and designed the experiments. Z.L. performed the majority of the experiments. R.Ki., V.T., G.M., G.B.C., A.F., B.M., and B.E. performed additional experiments. Z.L., R.Ki., and R.Kh. analyzed most of the experimental data. K.D., P.R., and B.E. helped with analyzing the results. V.T., A.B.O., and D.S. analyzed the sequencing data. Z.L. and R.Kh. wrote the manuscript with help from all authors. All the authors contributed to the integration and discussion of the results.

