## Supplement for "The ER cargo receptor SURF4 facilitates efficient erythropoietin secretion"

Fig. S1

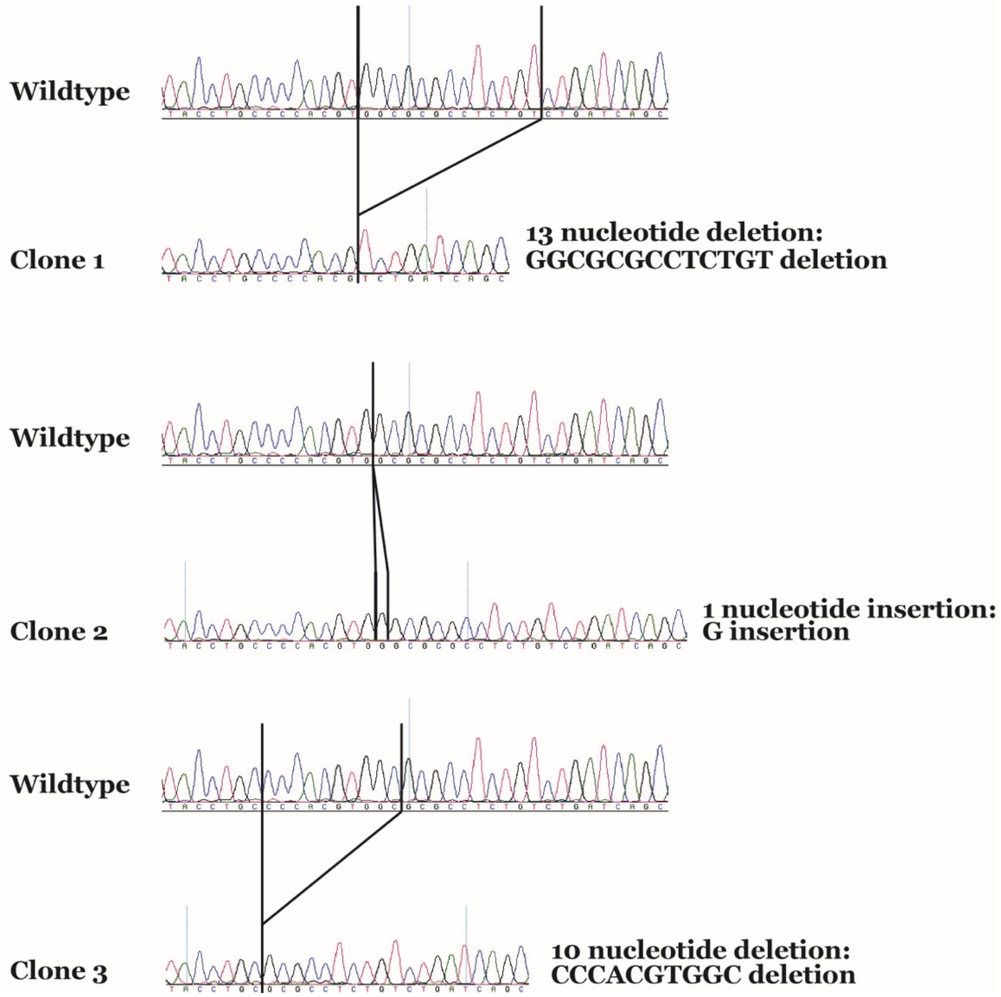


**Fig. S1. Generation of clonal SURF4 deficient reporter cells.** A *SURF4* targeting sgRNA was cloned into PX459 and transiently expressed in the reporter cell line. Genomic DNA was extracted and amplified. Three clonal SURF4 deficient cell lines (with frameshift mutations) were generated.

Fig. S2

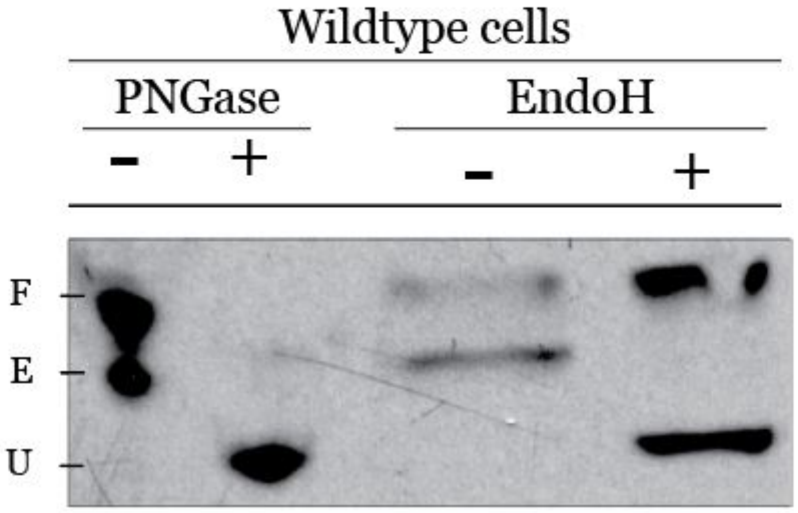


**Fig. S2. EPO glycosylation.** Cell lysates from wildtype cells expressing EPO-eGFP were collected and treated with EndoH, PNGase, or was left untreated. F = fully glycosylated EPO (post-Golgi from of EPO); E = ER form of EPO (endoH sensitive); U = unglycosylated EPO.

| Non-targeting gRNAs | sgRNA seqeuence 5'->3' |
|---|---|
| Non-targeting g1 | GTTCATTTCCAAGTCCGCTG |
| Non-targeting g2 | CGTGTGTGGGTAAACGGAAA |
| Non-targeting g3 | GTATTACTGATATTGGTGGG |
| Non-targeting g4 | TCATGCTTGCTTGGGCAAAA |

| SURF4 exon 2 targeting gRNAs | sgRNA seqeuence 5'->3' |
|---|---|
| SURF4 exon 2 g1 | TCAGACAGAGGCGCGCCACG |
| SURF4 exon 2 g2 | CAGGTAGCCGCAGTTCCAGG |

| SURF4 exon 2 sequencing primers | Primer seqeuence 5'->3' |
|---|---|
| Exon 2 forward | TCTGTTCCTCACACACCCCGCCC |
| Exon 2 reverse | ACTCACTCAGCTGTCCCAGCAAG |

| Other control sgRNAs used | sgRNA seqeuence 5'->3' |
|---|---|
| BCL11A g1 | TGAACCAGACCACGGCCCGT |
| BCL11A g2 | GCATCCAATCCCGTGGAGGT |
| MPL g1 | TGCCAGCAAGGAGACATCTA |
| MPL g2 | GTTCGGGAGAAACACTTCAG |
| SERPINA1 | CAATGCCGTCTTCTGTCTCG |
| EPO | GCCCAGAGGGAGCGACAGCA |

**Table S1. sgRNA and primer sequences.**
